# Embryonic development of *C. elegans* sense organs

**DOI:** 10.1101/2025.06.04.657903

**Authors:** Leland R. Wexler, Irina Kolotuev, Maxwell G. Heiman

## Abstract

*C. elegans* sense organs provide a powerful model for understanding how different cell types interact to assemble a functional organ. Each sense organ is composed of two glial cells, called the sheath and socket, and one or more neurons. A major challenge in studying their development has been the lack of methods to directly observe these structures in the embryo. Here, we mine a recently published high-resolution ultrastructural dataset of a comma-stage embryo that provides an untapped resource for visualizing early developmental events. From this dataset we reconstructed all head sense organs (two amphid (AM), four cephalic (CEP), six inner labial (IL), four outer labial quadrant (OLQ), and two outer labial lateral (OLL)). Symmetric sense organs were at different stages of morphogenesis, allowing us to infer developmental steps by which they form. First, we found that the sheath glial cell begins wrapping its partner neurons at the distal tip of the dendrites where it self-fuses into a seamless tube and then “zippers” down the dendrite. In many cases, sheath glia wrap the progenitors of partner neurons prior to their terminal division. After sheath wrapping has begun, the socket glia wraps the sheath glia circumferentially before presumably elongating to form the mature sheath-socket channel. We also observed transient interactions not found in the mature animal, such as amphid sheath glia wrapping the AUA neuron, that may reflect ancestral relationships. This study demonstrates the value of large public EM datasets that can be mined for new insights, and sheds light on how neurons and glia undergo coordinated morphogenesis.

## INTRODUCTION

Fifty years ago, two landmark papers in the *Journal of Comparative Neurology* described the three-dimensional structure of the anterior sensory anatomy of *C. elegans* – the first portion of its nervous system to be completely reconstructed (Ward et al., 1975; Ware et al., 1975). The use of serial section transmission electron microscopy to map every connection among sensory neurons and glia proved to be the first step in elucidating the total wiring diagram of the animal over the next decade. As the authors foresaw at the time, “This would open the way to understanding how the nematode’s behavior could be generated by the nervous structure” (Ward et al., 1975). Their studies also opened many new questions about development. They noted that, “Although individual worms are not precise replicas of each other down to the finest details they are remarkably exact copies” (Ward et al., 1975) and raised the question of what kind of developmental mechanism “produces a stereotyped nervous system with nearly 100% reproducibility” (Ware et al., 1975). Achieving the temporal and spatial imaging resolution required to answer this question has posed a major challenge. Thus, despite enormous progress, understanding how sensory neurons and glia accurately assemble into sense organs remains a mystery.

The sense organs in the head of *C. elegans* are each composed of two glial cells – called the sheath and socket glial cells – and one or more ciliated sensory neurons (reviewed in Singhvi and Shaham, 2019; Heiman and Bülow, 2024). In the mature animal, the sheath glial cell extends a process that wraps the ciliated dendrite endings in a seamless tube with ring-shaped adherens junctions around each dendrite (Ward et al., 1975; Doroquez et al., 2014; Low et al., 2019). The socket glial cell extends a process that is attached to both the sheath and hypodermis (skin) by adherens junctions. The socket forms an additional self-junction to create a tube-shaped channel, continuous with that of the sheath, through which the neuronal sensory cilia protrude. One of the major challenges of studying sense organ development is the lack of tools available to observe the formation of these structures in the embryo. The neurons and glia are produced during a burst of terminal cell divisions from ∼280-410 minutes post first cell cleavage (pfc), corresponding to a key morphogenetic period as the embryo progresses from a ball of cells to the “comma” stage (Sulston et al., 1983; Poole et al., 2024). While a powerful toolkit of cell-type-specific fluorescent markers is available to visualize individual sensory neurons and many glial cell types in larvae and adults (Gilleland et al., 2015; Fung et al., 2020), far fewer markers are known that are expressed at this stage in specific neurons and even fewer for specific glial subtypes. The paucity of early markers combined with the rapid twitching of the embryo after comma stage and insufficient spatial resolution to discern individual cell-cell contacts make live imaging of these early developmental events challenging.

Recently, Santella et al. published a series of high-resolution datasets obtained by focused ion beam scanning electron microscopy (FIB-SEM) and array tomography (AT-SEM) of entire embryos at four stages of development, combined with fluorescence-based cell lineage tracing to enable more accurate annotation of individual cells (Santella et al., 2022). These datasets provide an untapped resource for visualizing early developmental events that we previously lacked the tools to observe in such detail. Here we use one of these datasets to examine the sense organs in the head comprised of the two amphid (AM), four cephalic (CEP), six inner labial (IL), four outer labial quadrant (OLQ), and two outer labial lateral (OLL) sense organs at this snapshot of embryonic development. Among these, the amphid has been the most well studied, containing most of the ciliated sensory neurons in the head, while very little is known about the development of the other sense organs.

Focusing on the comma stage embryo dataset, we identified the glial and neuronal cells of all the sense organs through their stereotyped anatomy combined with the Santella et al. cell identity predictions, refining the single cell annotation of this embryo. We found the sense organs in intermediate stages of development, providing a picture of the dynamics of sense organ assembly. Based on our observations, we infer several steps through which the sense organ structure is formed: we propose that the sheath wraps the distal dendrite tips, fuses to itself, and then zippers up in a distal-to-proximal manner; meanwhile the socket initially wraps the sheath circumferentially before telescoping to the mature tube structure. We also identify previously undescribed transient interactions between non-ensheathed neurons and sheath glia cells. Our approach underscores the value of publicly available large EM datasets that can be repeatedly mined by investigators seeking to address different biological questions.

## RESULTS

### Cell identification and embryo timing

We first sought to identify all neurons and glial cells in the sense organs. Santella et al. used an algorithm to predict the identity of most cells by comparing their observed nucleus positions with known configurations of nuclei obtained from time-lapse fluorescence imaging of live embryos, an approach that was estimated to be ∼71-78% accurate (Santella et al., 2022). To unambiguously identify the cells in each sense organ, we first identified the 36 glial cells in the head and then used them to assign sensory neuron identities. The glia are comprised of paired sheath and socket glial cells for each of the 18 sense organs: two amphids, six inner labial (IL), six outer labial (four quadrant OLQ; two lateral OLL), and four cephalic (CEP) (Ward et al., 1975; Doroquez et al., 2014). The sheath and socket glial cell processes are easily distinguished from those of sensory neurons because they are substantially wider and can often be seen wrapping the neuron endings. The sense organ positions are highly stereotyped relative to each other and relative to the overall anatomy of the head. These positions, combined with the cell identity predictions made by Santella et al., allowed us to identify the glial cells of all sense organs. We then assigned sensory neuron identities based on their association with specific partner glial cells. By reconstructing the sheath glial cells and their associated neurons we found that the sense organs in our reconstruction are arranged symmetrically in their appropriate relative positions, supporting our identity assignments (Figure 1, Movie 1). Socket glia cells were identified but are not shown in Figure 1 or Movie 1 for clarity. Amphid morphology in this embryo was previously described (Santella et al., 2022) and is not included in Figure 1 or Movie 1.

**Figure 1.**
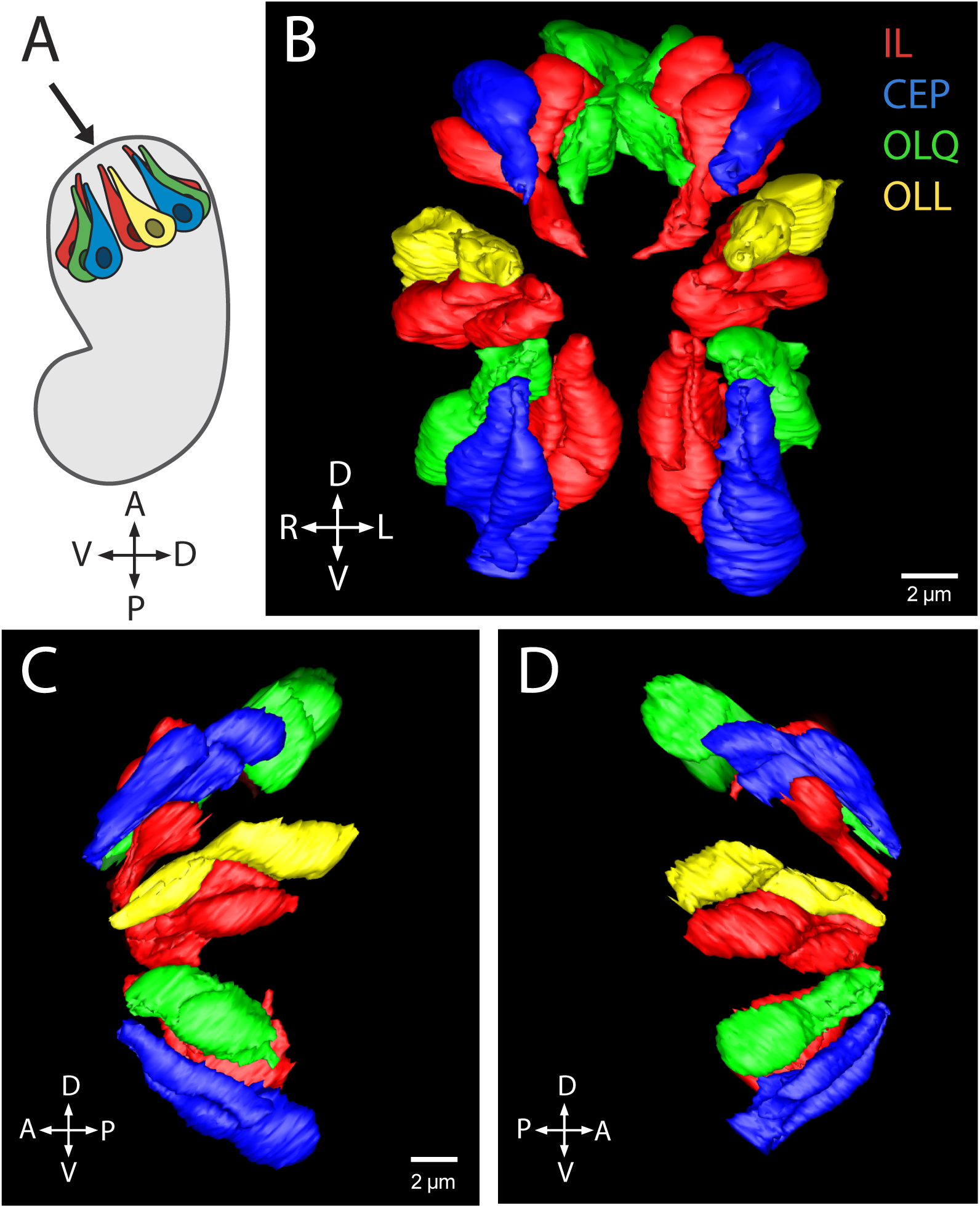
3D model of reconstructed IL, CEP, OLQ, and OLL sheath glia and their associated neurons (IL1 and IL2; CEP and CEM; OLQ; and OLL respectively). Sheath glia and neurons of the same class are in the same color (IL, red; CEP, blue; OLQ, green; OLL, yellow; amphids are not shown). (A) Cartoon of comma stage embryo with sense organs in the head. Arrow indicates view of panel B. (B) En face view. (C) Lateral view of left side. (D) Lateral view of right side.

Due to the rapid embryonic development of *C. elegans*, determining the age of this embryo is essential for understanding the developmental events occurring within it. Santella et al. estimated the embryo to be 345 min post first cell division (pfc) based on observed landmarks (Santella et al., 2022), however, we sought to further refine this temporal staging. While reconstructing the neurons associated with the sensory organs, we noticed that some of them appeared to be in the process of cell division. Whereas most nuclei exhibit a well-defined nuclear envelope with peripheral heterochromatin (Figure 2A), in some neurons and glia the nuclear envelope is fragmented, suggesting the cell is entering or exiting mitosis, and the heterochromatin is located centrally, suggesting chromosome condensation (Figure 2B). We aligned the CEP, IL1, OLQ, and OLL neurons by their established birth times (Sulston et al., 1983) and assessed which cells are in interphase and which are in or near cell division to estimate the age of this embryo (Figure 2C-R). We found that a cluster of cells born between 397-405 min pfc – IL1L, CEPVL, OLQDL, OLQDR, and OLLR – appear in or near cell division. All of these cells have centrally located heterochromatin and all but CEPVL have discontinuous nuclear envelopes (Figure 2G-K, S). Most of the neurons born after 405 min pfc are in interphase with two exceptions; IL1VR, born at 410 min pfc, is in mitosis and OLQVL, born at 415 min pfc, is likely near cell division, with centrally located heterochromatin but an intact nuclear envelope (Figure 2L-S). While the timing of cell divisions in the embryo is tightly regulated, there is a small degree of variability for each cell division that can accumulate in a lineage over time, and that may account for these discrepancies (Bao et al., 2008; Richards et al., 2013; Natesan et al., 2023). It is mostly likely that these cells in interphase are parent cells that have yet to begin their final cell division, although we cannot exclude the possibility that some may have already undergone their terminal cell division earlier than expected due to variability in cell division time. IL1DL and IL1DR, born at 390 min pfc, appear to be in interphase suggesting they have most likely already completed their terminal cell divisions. Taken together, these data bring our best estimate of the embryo age to ∼395 min pfc. At this timepoint the CEPD parent cells would have undergone their terminal divisions into CEPD and URX (380 min pfc) (Sulston et al., 1983). Consistent with this assessment, we identified a cell adjacent to CEP that is consistent with URX (Figure 2C-D), supporting our estimate of the embryo age as ∼395 min pfc.

**Figure 2.**
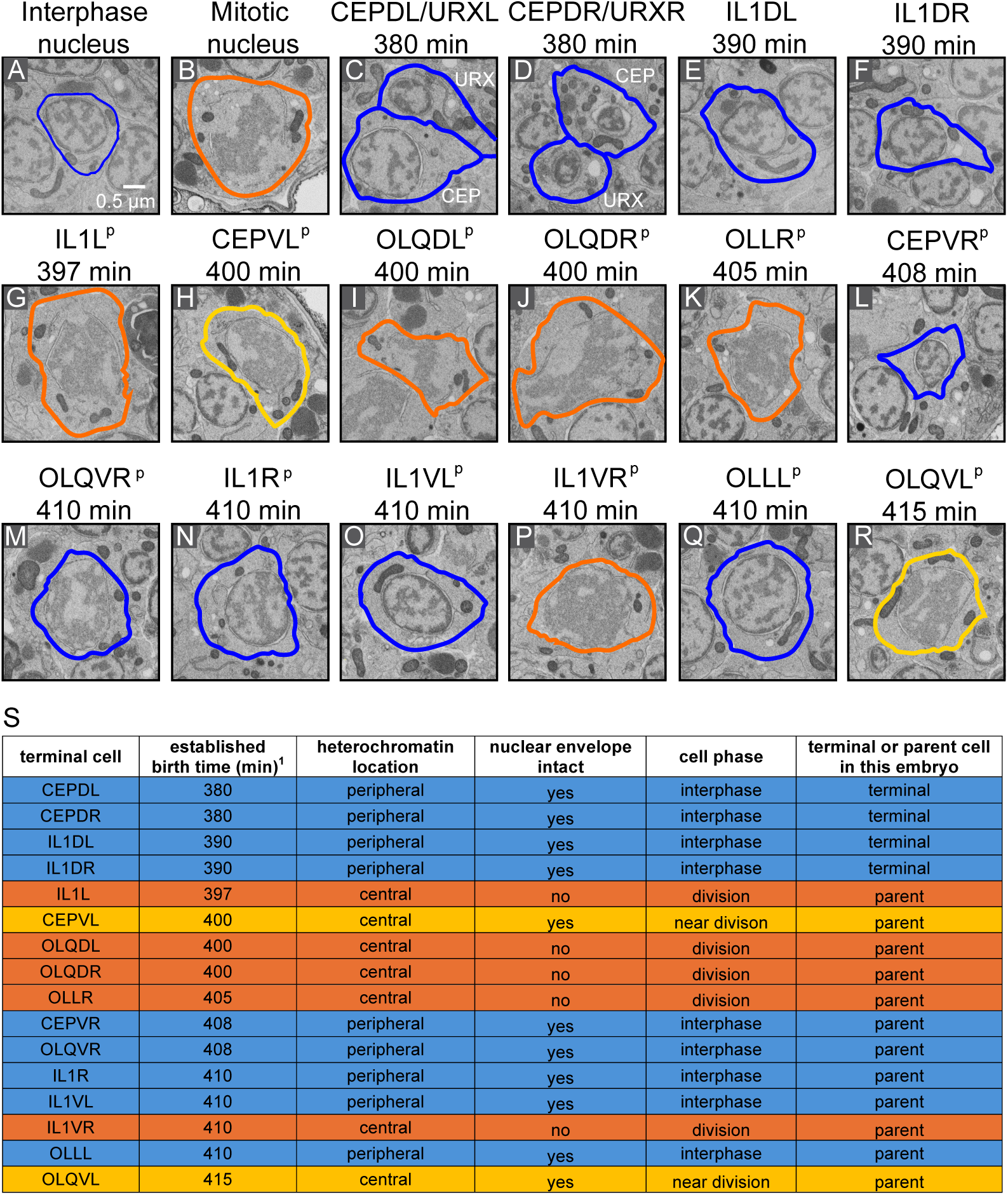
Embryo age was determined by which cells are in mitosis. (A) EM section of a cell in interphase (IL2DL) showing intact nuclear envelope and peripherally located heterochromatin. (B) EM section of a cell in mitosis (unidentified cell) showing discontinuous nuclear envelope and centrally located heterochromatin. (D-R) EM sections of CEP, IL1, OLQ, and OLL neurons ordered by established birth time (minutes post first cell division). ^p^ Indicates cell is a parent cell, for example IL1L^P^ is AB.alapaappa, which will divide to produce IL1L and a cell death. Blue outline indicates cell is in interphase (intact nuclear envelope, peripheral heterochromatin). Orange outline indicates cell is in mitosis (discontinuous nuclear envelope, central heterochromatin). Yellow outline indicates cell is likely to be entering or exiting mitosis (intact nuclear envelope but central heterochromatin). (S) Table summarizing data in A-R. Blue shading indicates cell is observed to be in interphase in this embryo. Orange shading indicates cell is observed to be in mitosis in this embryo. Yellow shading indicates cell is likely entering or exiting mitosis in this embryo. ^1^Sulston et al., 1983.

Comparing our identity assignments for these 36 glial cells and their 26 (non-amphid) associated sensory neurons to the predictions of Santella et al., we find that our assignments agree for 79% of cells overall, with 83% agreement for glial cells and 73% agreement for neurons (Table 1). By comparison, Santella et al. predicted the accuracy of their model to be 71-78% overall and 69-78% for head neurons (Santella et al., 2022). We note that the actual prediction accuracy for head neurons is likely higher as we find that all of the predicted amphid neurons are associated with the amphid sheath glial cell, confirming their identities as amphid neurons; however, as we did not attempt to confirm their individual identities through cell position, we did not include these in our calculations. Further, the algorithm used in Santella et al. sometimes assigned the same identity to two different cells (CEMDL, OLLshL, CEPDR, IL2DR) and in all cases our reconstruction supports one of the two assignments (Table 1). Overall, our findings based on reconstruction of sense organ anatomy confirm the accuracy of the Santella et al. model based on nucleus position within at least the range of accuracy they predicted. With the age of this embryo and cell identities established we next began to examine the development of the sense organs at this point in embryonic development.

**Table 1.** Comparison of this study’s single-cell identification to previous predictions (Santella et al., 2022). Green shading indicates same prediction. Yellow indicates different prediction. The prediction method by Santella et al. can result in two cells being assigned the same identity: ^a^Two cells assigned as CEMDL. ^b^Two cells assigned as OLLshL. ^c^Two cells assigned as CEPDR. ^d^Two cells assigned as IL2DR.

| Glia |  |  |  |
| --- | --- | --- | --- |
| This study | Santella et al. |  |  |
| This study | Santella et al. | This study | Santella et al. |
| AMshL | AMshL | AMsoL | AMsoL |
| AMshR | AMshR | AMsoR | AMsoR |
| CEPshDL | CEPshDL | CEPsoDL | CEMDL<sup>a</sup> |
| CEPshDR | CEPshDR | CEPsoDR | CEMDR |
| CEPshVL | CEPshVL | CEPsoVL | CEPsoVL |
| CEPshVR | CEPshVR | CEPsoVR | CEPsoVR |
| ILshDL | ILshDL | ILsoDL | ILsoDL |
| ILshDR | ILshDR | ILsoDR | ILsoDR |
| ILshL | ILshL | ILsoL | ILsoL |
| ILshR | no prediction | ILsoR | ILsoR |
| ILshVL | ILshVL | ILsoVL | ILsoVL |
| ILshVR | ILshVR | ILsoVR | ILsoVR |
| OLQshDL | OLQshDL | OLQsoDL | hyp3 |
| OLQshDR | OLQshDR | OLQsoDR | OLQsoDR |
| OLQshVL | OLQshVL | OLQsoVL | OLLshL<sup>b</sup> |
| OLQshDR | OLQshDR | OLQsoVR | OLQsoVR sister |
| OLLshL | OLLshL<sup>b</sup> | OLLsoL | OLLsoL |
| OLLshR | OLLshR | OLLsoR | OLLsoR |
| Neurons |  |  |  |
| This study | Santella et al. |  |  |
| This study | Santella et al. | This study | Santella et al. |
| CEPDL | no prediction | CEMDL | CEMDL<sup>a</sup> |
| CEPDR | CEPDR<sup>c</sup> | CEMDR | CEPDR<sup>c</sup> |
| CEPVL parent | CEPVL parent | CEMVL | CEMVL |
| CEPVR parent | CEPVR sister | CEMVR | CEMVR |
| IL1DL | URADL | IL2DL | IL2DL |
| IL1DR | IL2DR<sup>d</sup> | IL2DR | IL2DR<sup>d</sup> |
| IL1L parent | IL1L parent | IL2L | IL2L |
| IL1R parent | IL1R parent | IL2R | IL2R |
| IL1VL parent | IL1VL parent | IL2VL | IL2VL |
| IL1VR parent | IL1VR parent | IL2VR | IL2VR |
| OLQDL parent | OLQDL parent | OLQVR parent | OLQVR parent |
| OLQDR parent | OLQDR sister | OLLL parent | OLLL parent |
| OLQVL parent | OLQVL parent | OLLR parent | OLLR |

### Sheath glia begin ensheathing their neuronal partners at the distal tip

Most sensory neurons in *C. elegans* have a ciliated dendrite ending that is ensheathed by a specific sheath glial cell partner. Glial ensheathment of dendrites is a complex developmental process that results in the sheath glial cell forming a seamless tube around the dendrite ending, the dynamics of which are poorly understood. We considered two possible models: 1) the sheath cell wraps around the dendrite endings and fuses back to itself to create a tube by an extracellular fusion event, or 2) dendrites “tunnel” through the sheath glial cell, which hollows out a lumenal channel by intracellular fusion events. Previous analysis of the two amphid sense organs in this embryo suggested the first model is correct (Santella et al., 2022). Our reconstruction of the remaining 16 sense organs in the head further supports this conclusion and adds additional details. Importantly, because the sense organs we examined are at slightly different stages of morphogenesis, they allow us to infer some of the dynamics of this process even within this static snapshot of development.

Sheath cells appear to begin ensheathing their partner neurons at the very distal tip of the nascent dendrites. This is captured most clearly in the left OLL sense organ, with the distal tip of the OLLL parent cell being the only fully wrapped cross-section of the dendrite (Figure 3A-D; Movie 2). The four CEP sense organs are caught at different stages, providing a view of the progression of ensheathment. The CEPVR parent neuron is embedded in a surface depression of the sheath glial cell, but the sheath glial cell has not yet fully wrapped the neuron (Figure 4M-P; Movie 3). The CEPVL parent appears to be almost fully ensheathed at the distal tip of the dendrite, but the sheath glial cell has not yet fused into a tube (Figure 4I-L; Movie 3). Both dorsal CEP neurons are fully ensheathed along a large extent of the dendrite, possibly representing a completed wrap (Figure 4A-H; Movie 3). The progression of ensheathment is more difficult to see for the other sense organs. ILDL, ILDR, and ILVR have mostly or fully wrapped dendrites at the distal tips, with various degrees of ensheathment proximally down the dendrite, while ILL, ILR, and ILVL are not yet fully wrapped at the distal tip, although the IL1 neurons appear embedded in the sheath glial cell in these sense organs (Figure 5; Movie 4). None of the OLQ sense organs have fully ensheathed dendrites at this stage, but OLQs are deeply embedded in the sheath glial cells and it is clear the sheath glial cell has begun to project around the tip of OLQVL (Figure 6; Movie 5).

**Figure 3.**
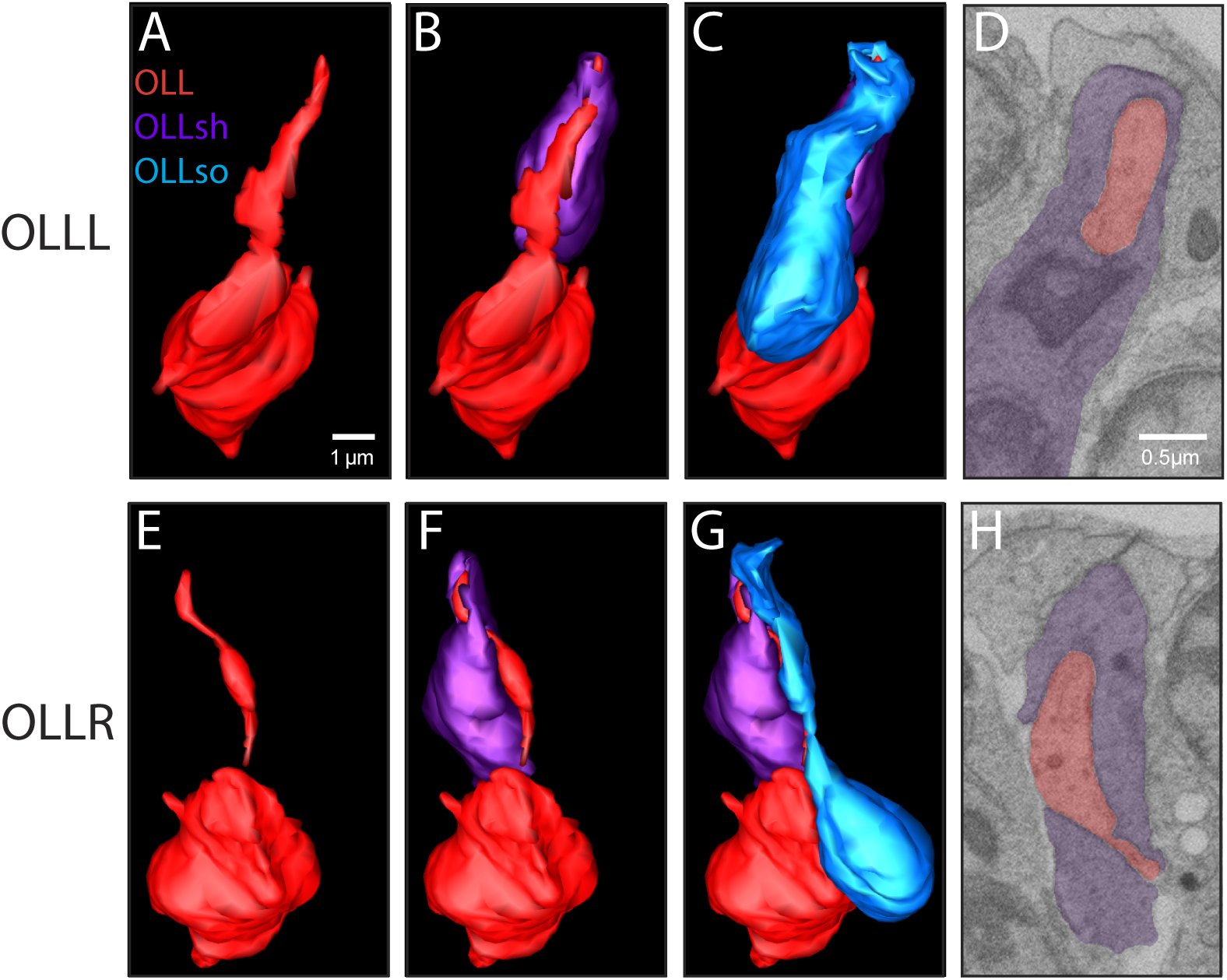
OLL sense organs. (A-C) 3D model of reconstructed OLLL neuron, OLLshL sheath glia, and OLLsoL socket glia. (D) EM section of OLLL and OLLshL. (E-G) 3D model of reconstructed OLLR neuron, OLLshR sheath glia, and OLLsoR socket glia. (H) EM section of OLLR and OLLshR. OLL neuron is red, OLLsh is purple, and OLLso is blue.

**Figure 4.**
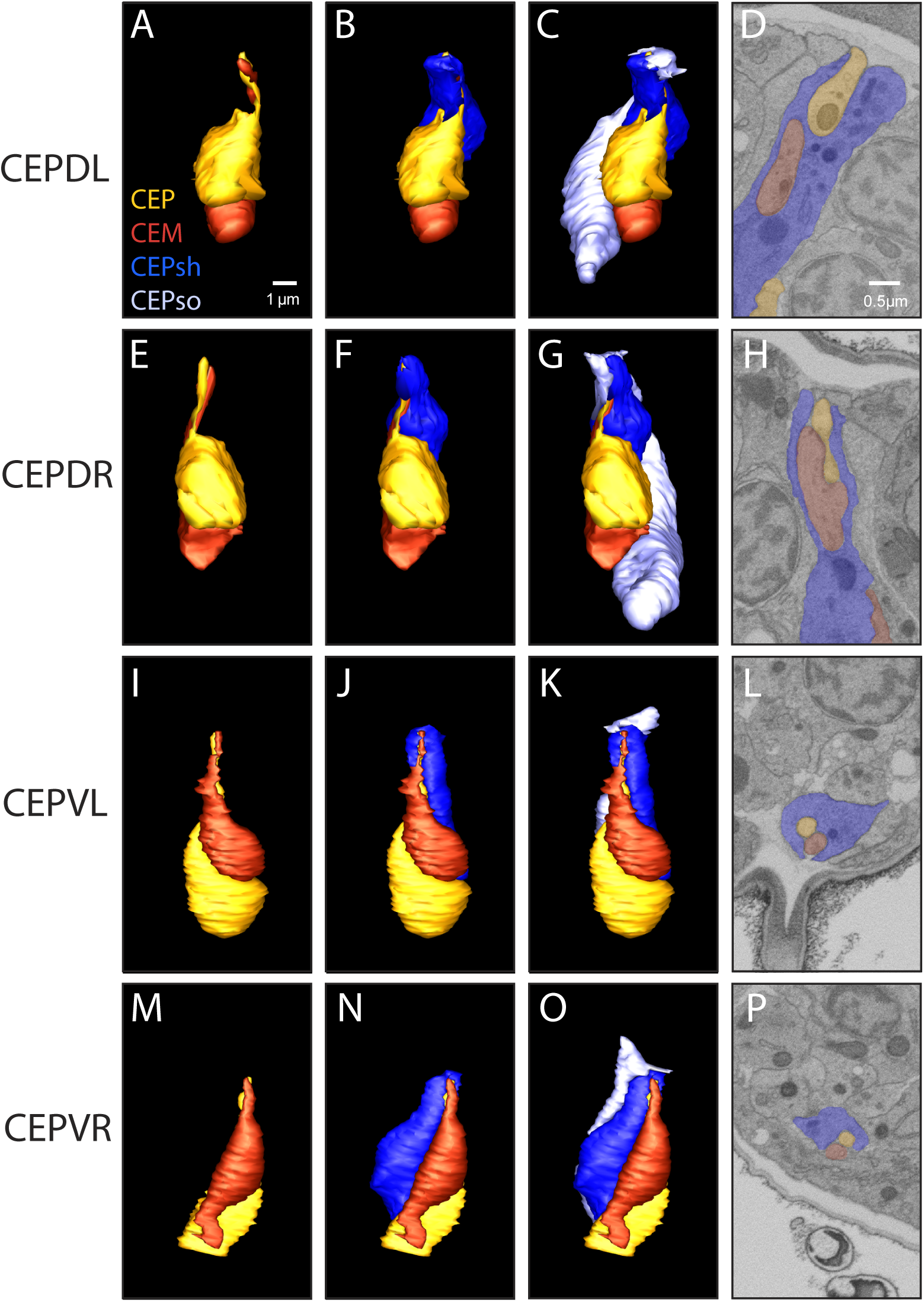
CEP sense organs. (A-C) 3D model of reconstructed CEPDL neuron, CEMDL neuron, CEPshDL sheath glia, and CEPsoDL glia. (D) EM section of CEPDL, CEMDL and CEPshDL. (E-G) 3D model of reconstructed CEPDR neuron, CEMDR neuron, CEPshDR sheath glia, and CEPsoDR socket glia. (H) EM section of CEPDR, CEMDR and CEPshDR. (I-K) 3D model of reconstructed CEPVL neuron, CEMVL neuron, CEPshVL sheath glia, and CEPsoVL socket glia. (L) EM section of CEPVL, CEMVL and CEPshVL. (M-O) 3D model of reconstructed CEPVR neuron, CEMVR neuron, CEPshVR sheath glia, and CEPsoVR socket glia. (P) EM section of CEPVR, CEMVR and CEPshVR. CEP neuron is yellow, CEM neuron is red, CEPsh is blue, and CEPso is lavender.

**Figure 5.**
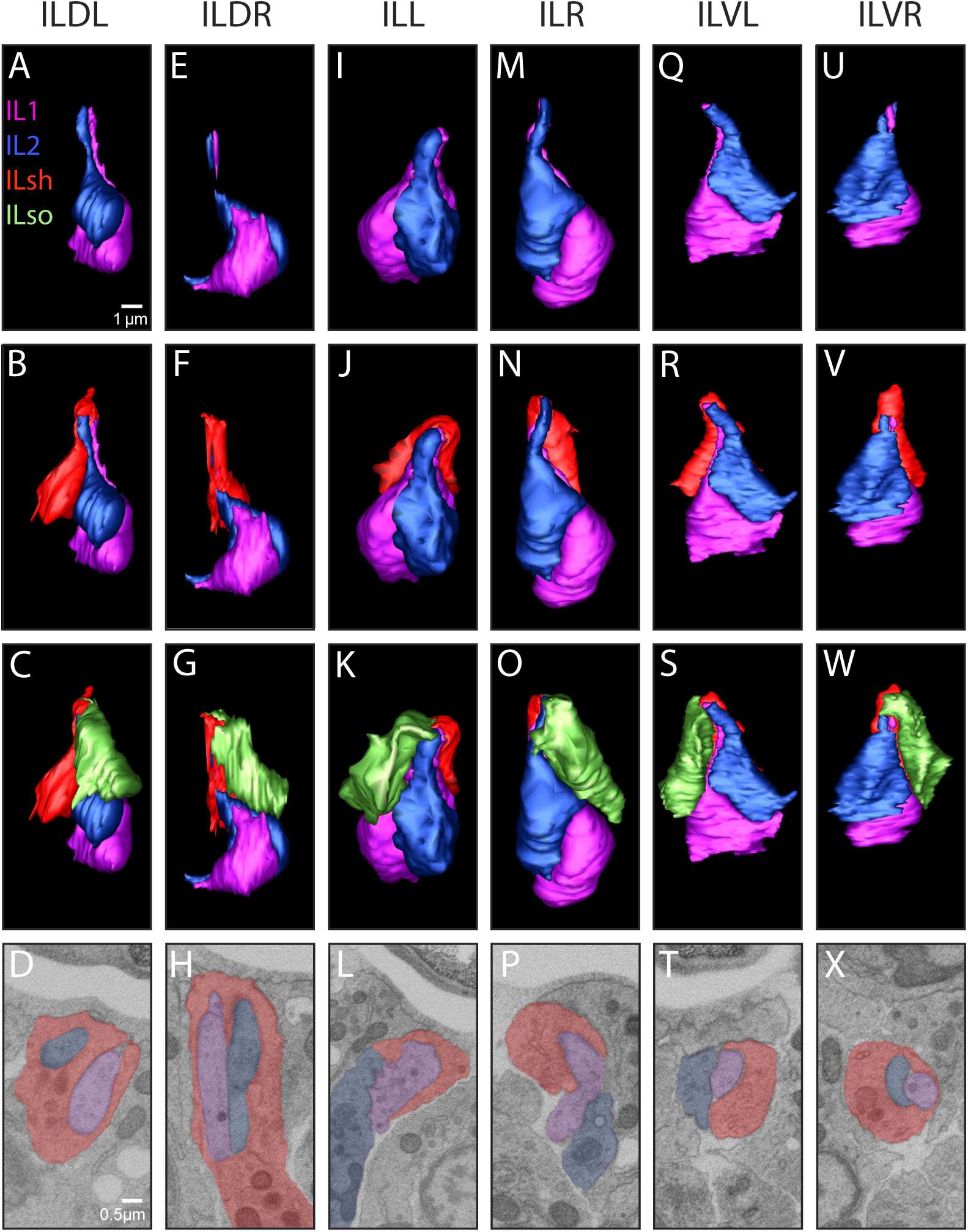
IL sense organs. (A-C) 3D model of reconstructed IL1DL neuron, IL2DL neuron, ILshDL sheath glia, and ILsoDL socket glia. (D) EM section of IL1DL, IL2DL and ILshDL. (E-G) 3D model of reconstructed IL1DR neuron, IL2DR neuron, ILshDR sheath glia, and ILsoDR socket glia. (H) EM section of IL1DR, IL2DR and ILshDR. (I-K) 3D model of reconstructed IL1L neuron, IL2L neuron, ILshL, and ILsoL. (L) EM section of IL1L, IL2L and ILshL. (M-O) 3D model of reconstructed IL1R neuron, IL2R neuron, ILshR sheath glia, and ILsoR socket glia. (P) EM section of IL1R, IL2R and ILshR. (Q-S) 3D model of reconstructed IL1VL neuron, IL2VL neuron, ILshVL sheath glia, and ILsoVL. (T) EM section of IL1VL, IL2VL and ILshVL. (U-W) 3D model of reconstructed IL1VR neuron, IL2VR neuron, ILshVR sheath glia, and ILsoVR socket glia. (X) EM section of IL1VR, IL2VR and ILshVR. IL1 neuron is pink, IL2 neuron is blue, ILsh is red, and ILso is green.

**Figure 6.**
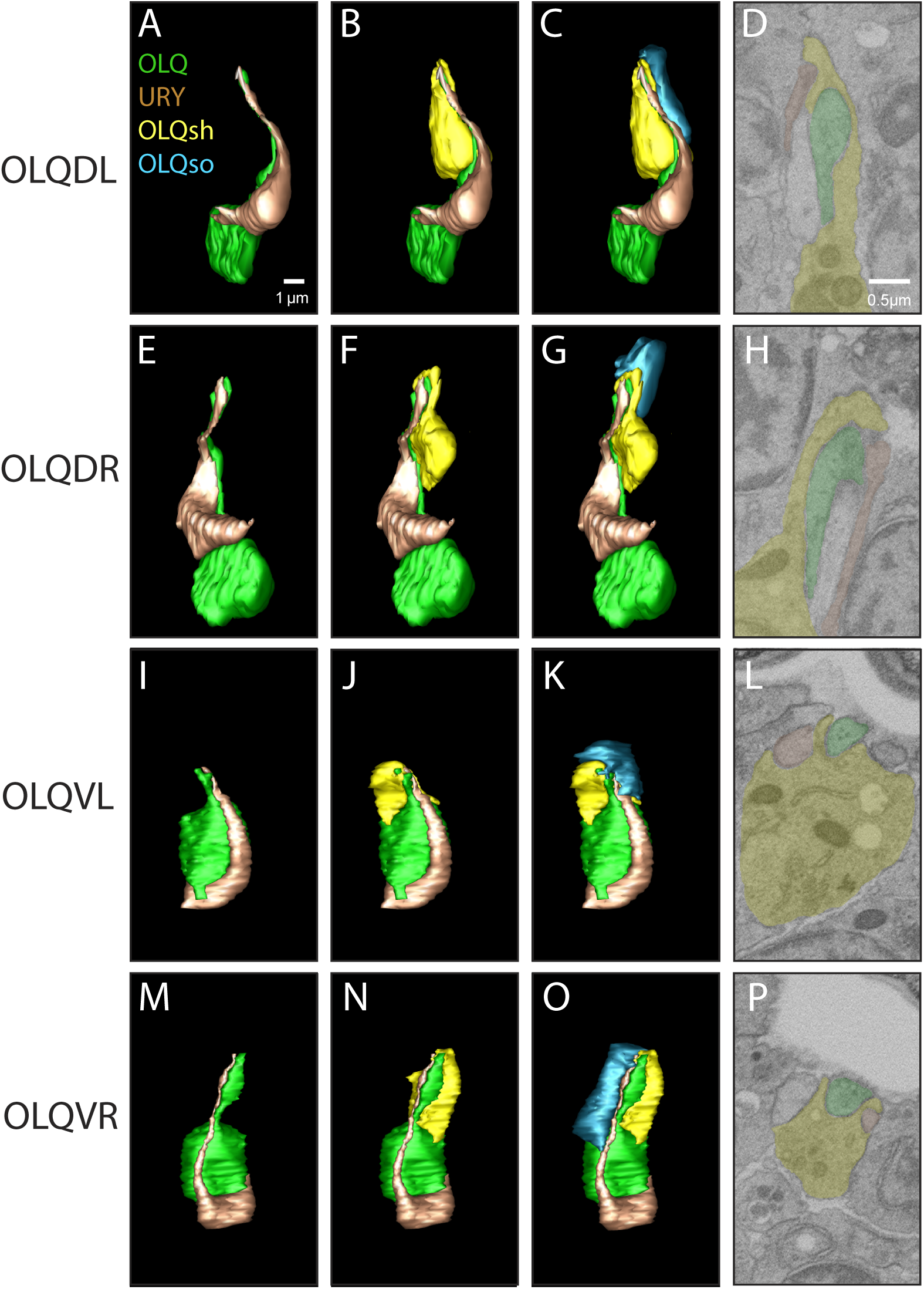
OLQ sense organs. (A-C) 3D model of reconstructed OLQDL neuron, URYDL neuron, OLQshDL sheath glia, and OLQsoDL socket glia. (D) EM section of OLQDL, URYDL and OLQshDL. (E-G) 3D model of reconstructed OLQDR neuron, URYDR neuron, OLQshDR sheath glia, and OLQsoDR socket glia. (H) EM section of OLQDR, URYDR and OLQshDR. (I-K) 3D model of reconstructed OLQVL neuron, URYVL neuron, OLQshVL sheath glia, and OLQsoVL socket glia. (L) EM section of OLQVL, URYVL and OLQshVL. (M-O) 3D model of reconstructed OLQVR neuron, URYVR neuron, OLQshVR sheath glia, and OLQsoVR socket glia. (P) EM section of OLQVR, URYVR and OLQshVR. OLQ neuron is green, URY neuron is brown, OLQsh is yellow, and OLQso is cyan.

It has been noted previously that amphid dendrites are ensheathed as a bundle during embryogenesis, and then later they become separated such that in the mature structure each dendrite individually penetrates the sheath glial cell (Oikonomou et al., 2011; Low et al., 2019; Heiman and Bülow, 2024). Further, the mature amphid sheath glial cell contains specialized lumenal compartments for the AFD, AWA, AWB, and AWC neurons, which do not enter the socket channel with the other amphid neurons. In this embryo we do not yet see this compartmentalization of the amphid (Figure 7; Movie 6), however, we observe different stages of compartment formation in the IL sense organs. Like the amphid neurons, the mature IL1 and IL2 neurons penetrate their sheath glial cell individually through separate openings that then merge into a single lumenal channel within the sheath. In this embryo the fully wrapped IL sense organs are at various stages of forming this separation. In ILVR, the neurons are fully ensheathed but no separation is yet seen between the IL1 and IL2 neurons (Figure 5U-X). In ILDR, the sheath glial cell appears to be forming a wedge between the IL1 and IL2 neurons starting at the distal tip of the neurons and working its way down, while in ILDL this separation is complete (Figure 5A-H).

**Figure 7.**
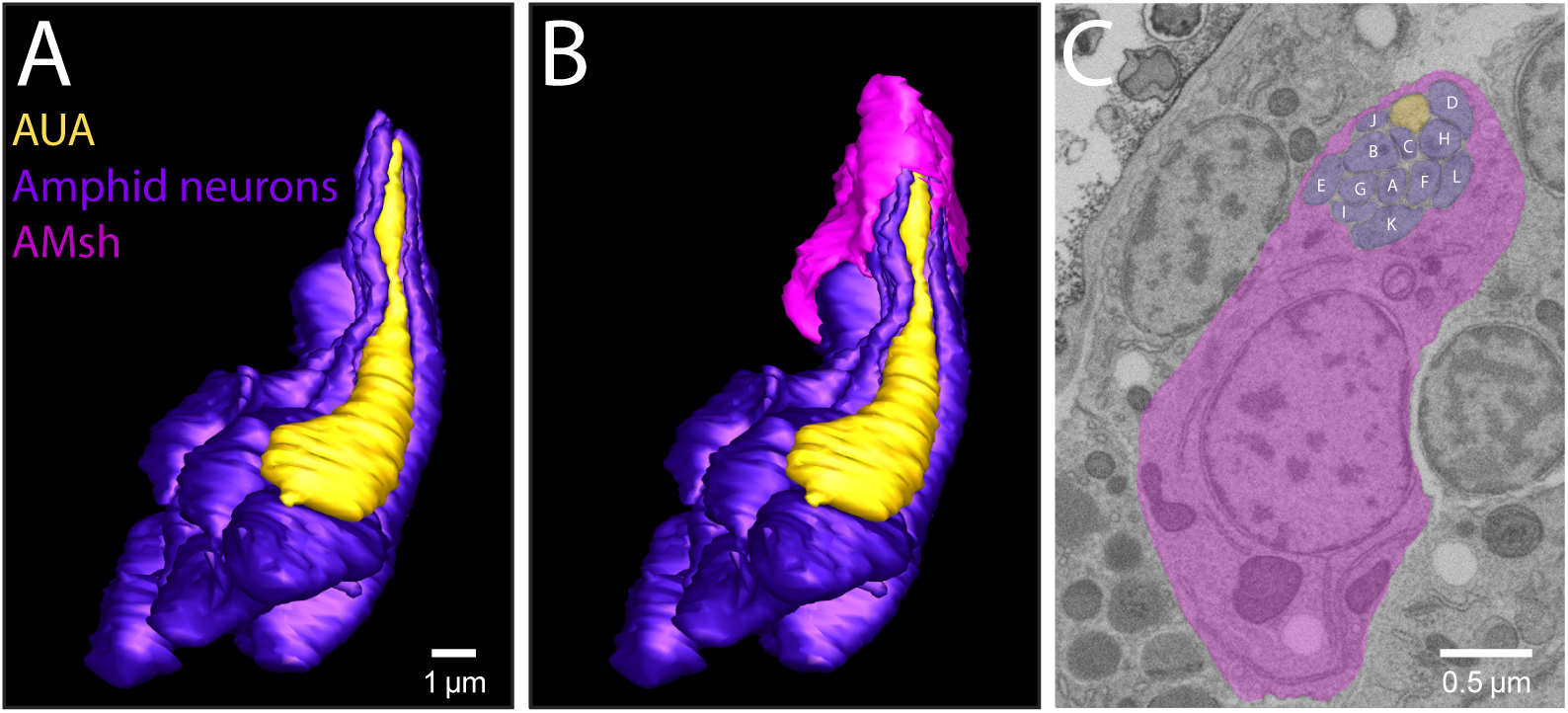
Left amphid sense organ. (A-B) 3D model of reconstructed left amphid neurons, left AUA neuron, and left AMsh sheath glia, (C) EM section of left amphid neurons, left AUA neuron, and AMshL. Amphid neurons are purple, AUA neuron is yellow, and AMsh is pink. A-L label predicted amphid neurons (AW<u>A</u>, AW<u>B</u>, AW<u>C</u>, AF<u>D</u>, AS<u>E</u>, AD<u>F</u>, AS<u>G</u>, AS<u>H</u>, ASI, AS<u>J</u>, AS<u>K</u>, AD<u>L</u>).

### Sheath glia begin ensheathing non-terminal cells

Many of the ensheathed cells we examined are neuronal parent cells that have yet to complete their final cell division (Figure 2). It has been previously suggested that neuronal parent cells can form dendrites as the CEPV parent has been found to have a dendritic process extending to the nose and, remarkably, to maintain that process during cell division (Shah et al., 2017). In this embryo, we found that the dendrites of many neuronal parent cells are either fully or partially ensheathed by their partner glial cells. This is most clearly seen in the OLL sense organs: the OLLL parent cell has not yet completed its final division yet its distal tip is fully ensheathed by the sheath glial cell, and the OLLR parent cell appears to be in the midst of cell division but also has its distal tip ensheathed (Figure 3; Movie 2). As another example, the CEPVL parent cell has not completed its final division but has also begun to be ensheathed (Figure 4I-L; Movie 3). Among the IL sense organs, all the IL2 neurons are terminal cells, but four of the IL1 cells (IL1L, IL1R, IL1VL and IL1VR) are parent cells. The IL1VR parent cell is an interesting example as it is in cell division and is fully ensheathed suggesting the glial cell wrapped the distal ending before this parent cell began mitosis (Figure 5U-X; Movie 4). Finally, none of the OLQ neurons have undergone their final division at this stage, yet ensheathment of their endings, while not completed, has begun (Figure 6; Movie 5).

Comparing sense organs of the same type, we noticed a correlation between the degree of ensheathment and whether the neurons were terminal cells or parent cells, suggesting the timing of glial morphogenesis is coordinated with the final neuronal cell division. For example, at this stage the dorsal CEP sense organs contain CEPD terminal neurons while the ventral CEP sense organs contain CEPV neuron parent cells. The dorsal CEPD terminal cells are extensively ensheathed by their sheath glial cells, while the ventral CEPV parent cells are not (Figure 4; Movie 3). Of the two ventral CEPV parent cells, the CEPVL parent cell is near cell division and has begun ensheathment at its distal tip, while the CEPVR parent cell has not begun mitosis and is at a much earlier stage of ensheathment (Figure 4I-P; Movie 3). This correlation is seen in the IL sense organs as well, with the dorsal IL1 and IL2 neurons having completed their final division and being fully ensheathed while the ILR and ILVL sense organs, containing IL1 parent cells, are not yet ensheathed at the tip (Figure 5; Movie 4). The IL1L and IL1VR parent cells are both in cell division; the ILL neurons are not yet ensheathed while the ILVR neurons are fully ensheathed (Figure 5; Movie 4), suggesting that ensheathment of the IL neurons occurs sometime during the terminal cell division of IL1. The timing of ensheathment could differ between sense organs, for example both the OLLL and OLLR parent cells are ensheathed at the distal tip (Figure 3; Movie 2) making the OLLL parent the only pre-mitotic parent cell to be fully ensheathed. For this reason, we have focused on comparisons between sense organs of the same class. The correlation in the timing of the terminal division and ensheathment, seen within the CEP and IL sense organ classes, raises the possibility that completing the final cell division helps to trigger full ensheathment.

### The CEM neuron is ensheathed prior to death

Interestingly, the male-specific CEM neuron is also wrapped by the CEP sheath glial cell (Figure 4; Movie 3). The CEM neurons are born in both sexes from the same cell lineages at ∼320 min pfc but undergo programmed cell death in hermaphrodites at ∼470 min pfc (Sulston et al., 1983) and survive only in males. This embryo was obtained from a hermaphrodite-only culture. While we cannot exclude the possibility that the embryo happened to be a spontaneous male, this observation suggests that CEM neurons are ensheathed after birth in both sexes, prior to undergoing hermaphrodite-specific cell death later in development. This observation is consistent with previous work showing that the hermaphrodite CEM neuron extends a dendrite prior to undergoing programmed cell death (Singhal and Shaham, 2017). How the channel of the CEP sense organs may be remodeled after the death of CEM to accommodate one neuron instead of two remains unknown.

### URY and AUA associate with the OLQ and amphid sense organs respectively

While examining the OLQ sense organ we observed that a second neuron in addition to the OLQ parent cell was closely associated with the OLQ sheath glial cell. This neuron is predicted to be URY (Santella et al., 2022) and its location is consistent with that prediction. In a mature animal, the four URY neurons (URYDL, URYDR, URYVL, URYVR) each project a non-ciliated dendrite to the tip of the nose, terminating in branched processes, positioned adjacent to the OLQ sheath glial cells and near the dorsal or ventral IL sheath glial cells (Doroquez et al., 2014). In this embryo, the URY neurons appear to be partially embedded in the OLQ sheath glia, which is most apparent in the OLQVL and OLQVR sense organs (Figure 6I-P; Movie 5). Unlike the OLQ parent cell, URY is not in the process of being ensheathed by the OLQ sheath (Figure 6; Movie 5). It is possible that the apparent interaction of URY with the OLQ sheath glial cell arises solely from proximity in the crowded environment of the embryo, however we have not observed this kind of partial embedding for other non-ensheathed dendrites. For example, a neuron predicted to be URA is also adjacent to the OLQ sheath glial cell and runs tightly alongside it but is not embedded (not shown). Notably, URY is the sister of the OLQ parent cell, raising the possibility that cell-surface proteins on the OLQ parent cell that allow it to be recognized by the OLQ sheath glial cell might also be present on the surface of URY.

We found an even more intimate and unexpected association between another neuron and the amphid sheath glial cell. In this embryo, the amphid sheath glial cell appears to have completely ensheathed all 12 amphid neurons; however, a thirteenth neuron, predicted to be AUA, is also ensheathed (Figure 7). The position of this neuron is consistent with the location of AUA and is adjacent to its sister cell, the predicted ASJ neuron. Although we did not fully reconstruct the right amphid sense organ, we found that it also includes an ensheathed AUA. In mature animals, the AUA dendrite projects approximately halfway along the amphid bundle, terminating just past the anterior pharyngeal bulb, far from the nose and not ensheathed by the amphid sheath glial cell (White et al., 1986). It was previously shown that AUA associates with the amphid in early morphogenetic (bean and comma stage) embryos (Fan et al., 2019). At this stage, the amphid neurons and glia form a multicellular rosette that also includes AUA and four other non-amphid neurons (Fan et al., 2019). The embryonic ensheathment of AUA suggests that AUA remains associated as part of the amphid for some time, possibly during much of dendrite extension. How and when it dissociates from the sheath glial cell is unknown. Notably, in some nematode species – including *Acrobeles complexus*, *Parastrongyloides trichosuri,* and *Strongyloides stercoralis –* the mature amphid contains a thirteenth ensheathed, ciliated neuron thought to be the homolog of AUA (Ashton et al., 1995; Bumbarger et al., 2009; Zhu et al., 2011). It is interesting to speculate that the embryonic association of AUA with the amphid sense organ, and possibly URY with the OLQ sense organ, may represent an ancestral state from which *C. elegans* has diverged.

### Socket glial cells circumferentially wrap sheath glial cells following neuron ensheathment

In a mature sense organ, the socket glial cell ending sits atop that of the sheath glia cell so that they form a continuous channel through which neuronal cilia protrude to reach the external environment. In this embryo, the socket glial cells are in various stages of morphogenesis that suggest they wrap around the circumference of the sheath glial cell before telescoping to form the mature channel. In some sense organs, such as ILL, ILR, ILVL, ILVR, OLQDL, OLQDR, and OLQVR, the process has not yet begun and the tip of the socket glial cell is positioned adjacent to the sheath glial cell but has not started to wrap around it (Figures 5, 6; Movies 4, 5). In ILDL, ILDR, OLQVL, and the CEP sense organs, intermediate stages of the process are seen with the socket glial cell beginning to wrap around the circumference of the sheath glial cell (Figures 4-6; Movies 3-5). Finally, in the OLL sense organs, this process appears nearly complete with a channel formed through which the OLL parent cell is visible (Figure 3; Movie 2). Interestingly, in all sense organs, the process of forming the socket glial cell channel seems to correlate with the extent of neuronal ensheathment by the sheath glial cell.

## DISCUSSION

This snapshot of embryonic development provides a window into the dynamics of morphogenesis as the sensory neurons, socket, and sheath glial cells form the sense organ structure. We find that ensheathment of neurons begins at the distal tip of the dendrite and works its way down, with the socket glial cell wrapping around the circumference of the sheath glial cell after neuronal ensheathment is mostly or fully complete (Figure 8). While these EM data show us how sense organ structures form, the molecular mechanisms initiating and guiding these developmental processes are poorly understood. Some proteins involved in forming or maintaining sense organ structure have previously been identified including the extracellular matrix proteins DYF-7 and DEX-1 (Heiman and Shaham, 2009; Low et al., 2019). Loss of DYF-7 or DEX-1 results in severely shortened amphid neurons and amphid sheath glial cells that fail to reach the nose, with the tips of the shortened dendrites still ensheathed by the sheath glial cell and separated from the amphid socket glial cell which remains attached to the hypodermis (Heiman and Shaham, 2009; Low et al., 2019). This phenotype fits with the order of development we see in this embryo, with the sheath glial cell and neurons first forming a morphological unit and the socket glial cell only later joining this structure, suggesting DYF-7 and DEX-1 may act in the time frame after neuron ensheathment and before or during attachment of the socket glial cell to the sheath glial cell. The CEP, OLQ, and IL sense organs are also affected by DYF-7 but to a much lesser extent, indicating some molecular differences in the development of different sense organs (Low et al., 2019).

**Figure 8.**
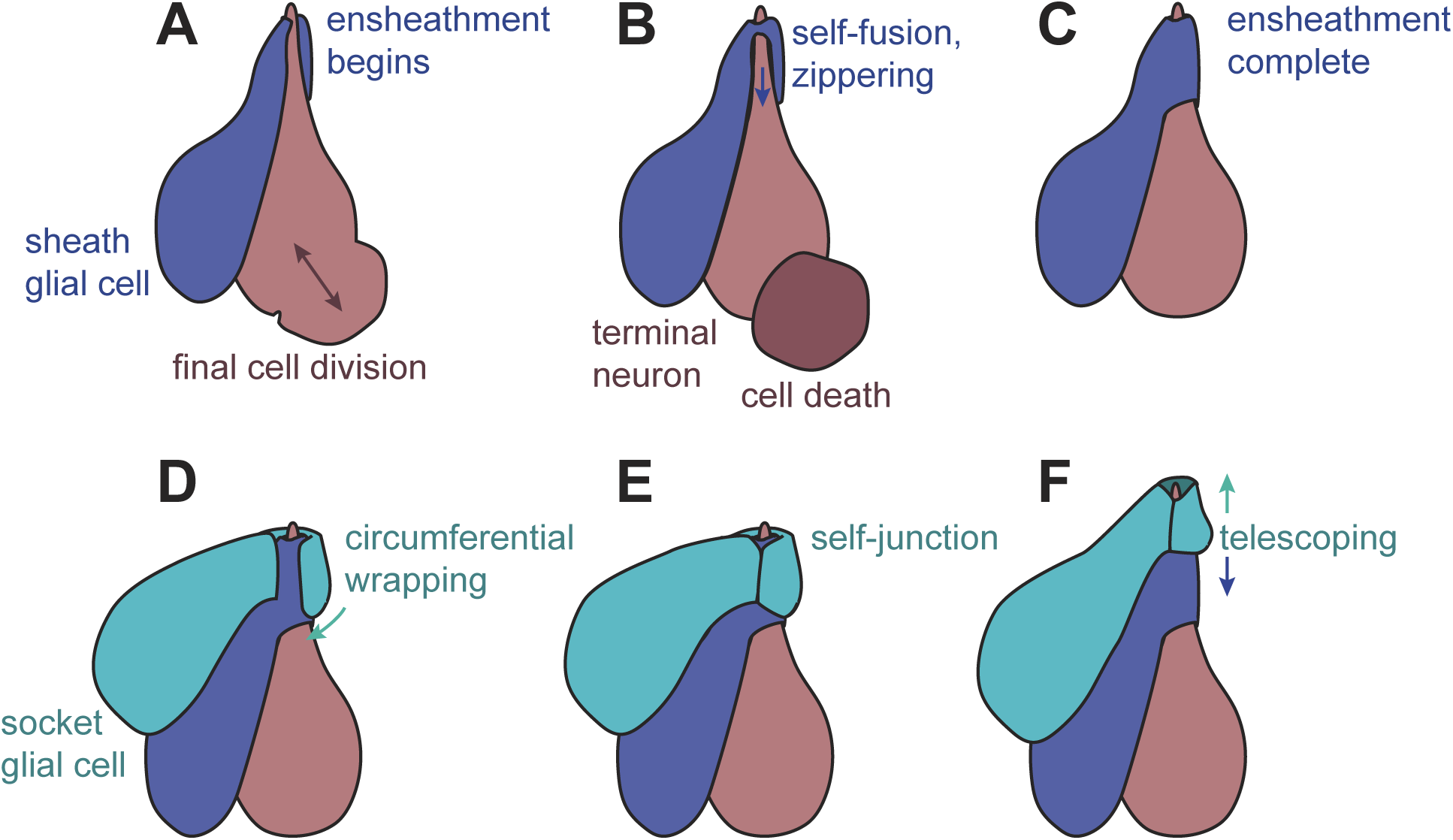
Proposed model of sense organ development. (A) Sheath glia begin wrapping the distal dendrite of neuron or neuronal progenitor as it completes mitosis. (B, C) Sheath glia undergoes self-fusion at the distal ending to form a seamless tube and then zippers proximally to complete ensheathment. (D) Socket glia wraps sheath circumferentially and (E, F) presumably then forms its mature self-junction and telescopes distally to complete the glial tube.

We show that sheath glial cells begin ensheathing neuronal parent cells, suggesting that they are able to recognize their partner cells before they are fully differentiated. For all the parent cells we examined, the final division results in a terminally differentiated neuron and a cell death. In these cases, division may occur at the soma such that the terminal neuron inherits the elongated dendrite and glial ensheathment of the parent cell (Figure 8). It is not clear how a cell can divide while maintaining its morphology and cell-cell attachments, but a similar event occurs in the male amphid socket glial cell as it undergoes a post-embryonic cell division to generate a neuron while maintaining its elongated morphology and tube-shaped ending (Sammut et al., 2015). By contrast, some parent cells may need to divide to produce two terminal neurons with long dendrites, such as CEPD and URX. In this embryo the CEPD/URX parent cells have already completed their final cell division (380 min pfc), but it would be intriguing to know if these parent cells are ensheathed prior to dividing to produce two terminal neurons with long dendrites, one of which (CEPD) is ensheathed while the other (URX) is not.

When comparing different sense organs of the same type, the correlation between the “age” of the sensory neuron (time since its final cell division) and the progress of ensheathment suggests that some timing cue may trigger ensheathment. One possible cue is the onset of cilia formation, which involves conversion of the centriole into a ciliary basal body and thus cannot occur until the final cell division is complete. Notably, mutations in *grdn-1* affect positioning of the centriole/basal body at dendrite endings as well as glial ensheathment, consistent with a mechanistic link between these events (Nechipurenko et al., 2016).

The associations between URY and the OLQ sheath glial cell, as well as AUA and the amphid sheath glial cell, raise many questions about how non-ensheathed neurons interact with sheath glial cells during development, and, in the case of URY, possibly in adults. Due to their shared lineage, URY and the OLQ parent cells may express the same cell-surface factors that lead to recognition by the OLQ sheath glial cell. The adjacency of URY to the OLQ sheath glial cell in adults suggests the possibility that a specialized neuron-glia interaction persists between these cells in the mature animal. For example, two sensory neurons, BAG and URX, are not ensheathed yet form specialized connections with the lateral IL socket glial cell (Cebul et al., 2019). These observations suggest that other such specific neuron-glia interactions may remain to be discovered or may occur transiently during development.

To our knowledge, the transient embryonic wrapping of AUA by the amphid sheath glial cell has not been previously described. The structure and function of amphid sensory neurons appears to be broadly conserved across species, and it has been proposed that AUA is homologous to amphid neurons in other nematode species. *Acrobeles complexus* has a thirteenth amphid neuron, called ASA, that may be the homolog of *C. elegans* AUA (Bumbarger et al., 2009). *A. complexus* ASA has a distinct morphology from other amphid neurons – it extends a dendrite to the nose and is ensheathed by the amphid sheath glial cell but has only a rudimentary cilium that terminates at the base of the amphid sheath glial channel (Bumbarger et al., 2009). A thirteenth amphid neuron has also been described in the parasitic nematode *Strongyloides stercoralis* and the related species *Parastrongyloides trichosuri* (Ashton et al., 1995; Zhu et al., 2011). Thus, it is possible that developmental interactions in the *C. elegans* embryo may reveal ancestral neuron-glia arrangements in nematode sense organs.

## METHODS

### Data source

All analysis was performed on the dataset “c_elegans_embryo_345min” obtained by FIB-SEM analysis of a 27.5 µm x 26.5 µm x 41.0 µm volume (1641 sections each of approximately 25 nm thickness) as previously described (Santella).

### Cell tracing and reconstruction

Cells were manually traced in IMOD as individual objects using the Model mode. All 3D graphical models were generated from these tracings in IMOD using the Model View mode (Kremer et al., 1996).

## ACKNOWLEDGEMENTS

We thank Anthony Santella, Zhirong Bao, and members of the Heiman lab for helpful discussions and comments.

